# Stereotyped cell lineage trees support robust development

**DOI:** 10.1101/2024.02.23.581522

**Authors:** Xiaoyu Zhang, Zizhang Li, Jingyu Chen, Wenjing Yang, Peng Wu, Feng Chen, Xingxing He, Xiaoshu Chen, Jian-Rong Yang

**Author notes:** Corresponding authors. (Yang JR). These authors contributed equally to this work (Xiaoyu Zhang, Zizhang Li).

## Abstract

Multicellular organisms must have robust development to ensure physiological stability in the face of environmental changes or perturbations. While various mechanisms contributing to developmental robustness have been identified at the subcellular level, those at the intercellular and tissue level remain largely unknown. Our study explores this question using an in vitro directed differentiation model of human embryonic stem cells (hESCs) into lung progenitor cells. Integrated analysis of single-cell transcriptomes and high-density cell lineage trees (CLTs) of the same colonies allowed a fine-resolution recapitulation of known cell types, as well as their differentiation hierarchies and developmental trajectories. Most importantly, we observed stable cell type compositions among many sub-CLTs across biological replicates. Systematic comparison among CLTs by a novel computational framework for CLT alignment suggests that stereotypical development extends beyond stable cell type composition to a degree of significant resemblance in sub-CLT topology. The existence of such sub-CLTs resembling each other not only deepens our understanding of developmental robustness by demonstrating the existence of a stereotyped program, but also suggests a novel perspective for the function of specific cell types in the context of stereotyped sub-CLTs.

## Introduction

Developmental robustness, also known as canalization^1^, refers to the phenomenon that biological development outcomes remain largely unchanged despite environmental or genetic perturbations^2,3^. In addition to being an essential feature of complex organisms, developmental robustness also has profound implications for evolution^4,5^ and disease^6^. Decades of studies have identified a variety of mechanisms that contribute to developmental robustness, including chaperone proteins^7^, microRNAs^8–10^, morphology-stabilizing genes^11,12^, feedback loops^13^, molecular redundancies^14^ and defect-buffering cellular plasticity^15^. While significant advances have been made at the molecular/intracellular level, other mechanisms that ensure robust development at the intercellular/tissue levels remain poorly understood. A couple examples include the nonlinear relationship between key regulators’ gene expression and embryonic structures^16^, and the robustness to cell death observed for determinative developmental cell lineages^17^.

The developmental process encompasses both the history of cell divisions and state transitions ^18,19^. It is thus possible to examine development, as well as its robustness, from two perspectives. In the first, cellular states, such as single-cell transcriptomes, were recorded during various developmental stages and used to construct a continuum of states known as an epigenetic landscape^20,21^ or state manifolds^18^. In the second, all cell divisions since the zygote or some progenitor cells can be recorded and used to construct a cell lineage tree (CLT)^22^. This CLT-based perspective, however, has been much less studied due to the difficulty in obtaining CLTs in complex organisms. Nonetheless, recent technological advancements in CLT reconstruction, particularly those utilizing genomic barcoding^19^, have led to new opportunities for joint analyses of these two perspectives. For example, scGESTALT simultaneously determined cell states by single-cell transcriptomics and the corresponding CLT via lineage barcodes^23^. Similar methods^18,19^ provide a combined view of single-cell states and CLTs, enabling CLT-based analyses of robustness for different developmental models.

One of the main manifestations of developmental robustness is the generation of adequate numbers of cells of various types in an appropriate cellular composition, especially when they work together as a functional unit. For example, the *Drosophila* peripheral nervous system contains thousands of identical mechanosensory bristles^24^, each consisting of exactly one hair cell, one socket cell, one sheath cell and one neuron^25^. Another well-known example is the functional unit of the endocrine pancreas, the islet, which has been shown in mice to consist predominantly (∼90%) of β cells at the core and □ and δ cells in the periphery^26^. To identify potential CLT characteristics that contributed to such a manifestation of developmental robustness, two CLT-based studies are particularly relevant. In the first, it was found that development of mammalian organs is preceded by significant mixing of multipotent progenitor cells^27^. Therefore, most organs have a polyclonal origin that ensures sufficient number of cells even some progenitors failed^27^. In the second, CLT of cortical development revealed stereotyped development giving rise to monophyletic clades of mixed cell types^28^. On the basis of these observations, we hypothesized that the combination of polyclonal origin and stereotyped development facilitates the robust development of adequate numbers of cells with an appropriate cellular composition. It is imperative to note that as our hypothesis revolves around the above-mentioned functional units, CLTs with sufficient resolution (fraction of cells sampled) are essential, otherwise stereotyped development cannot be detected with only <1% cells sampled from each functional unit. In addition, a high resolution CLT would also reveal how stereotyped development occurs, such as mitotic-coupling versus population-coupling development^18^ and whether epigenetic memory^29^ plays a role.

To this end, we obtained the single-cell transcriptomes and high density (capturing > 10% cells in the culture) CLTs of three *in vitro* cell cultures that mimic the *in vivo* development of human embryonic stem cells (hESCs) into lung progenitors ^30^. According to a joint analysis with *in vitro* cultures that retained stemness, single-cell transcriptomes were clearly separated into clusters of undifferentiated and various differentiated cell types, and the CLTs showed significant signals of divergence among subclones consistent with known sequential involvement of Bmp/TGF-β, Wnt and other endoderm differentiation related pathways. Multiple monophyletic groups of cells with highly stable cellular compositions were revealed by this CLT, providing direct support for our hypothesis of polyclonal stereotyped development. Furthermore, we found that some sub-CLTs with similar topological structures and terminal cell type compositions are significantly overrepresented, suggesting that at least some stereotyped development is driven by a mitotic coupling process.

Together, we demonstrate the existence of stereotyped lineage trees, a feature of CLTs that likely contributes to stable cellular composition and therefore developmental robustness.

## Results

### Reconstructing high-density cell lineage trees for directed differentiation of primordial lung progenitors

We aimed to determine the CLT of embryonic stem cells undergoing *in vitro* directed differentiation towards lung progenitors according to a well-established protocol recapitulating *in vivo* development^30^. This *in vitro* model of directed differentiation was chosen for several reasons. First, cells cultured in a small petri dish have a relatively homogenous environment, so that transcriptome divergence caused by environmental factors, or phylogeny-independent convergence due to niche-specific signals is unlikely. Second, the development trajectory of embryonic stem cells to the lung is well-known, such that the *in vitro* cell culture can be monitored to ensure that they closely mimic physiological situation. Indeed, our implementation of the protocol can reach the alveolar epithelial cells (AEC2s) fate after 20 days of directed differentiation. Third, *in vitro* culture allows us to induce Cas9 expression and therefore initiate the editing of the lineage barcode concurrently with the directed differentiation. Last but not least, it allows better control over the number of cells within the colony assayed for single-cell transcriptomes and CLTs. In particular, our cell culture begins with ∼10 hESCs and ends with ∼ 5,000 cells on day 10, of which a relatively high percentage can be captured in downstream experimental pipelines of 10x Chromium. The ten-day directed differentiation covers three critical phases of lung development, including definitive endoderm (DE), anterior foregut endoderm (AFE) and NKX2-1^+^ primordial lung progenitor (PLP)^30^ (**Figure 1A**).

**Figure 1.**
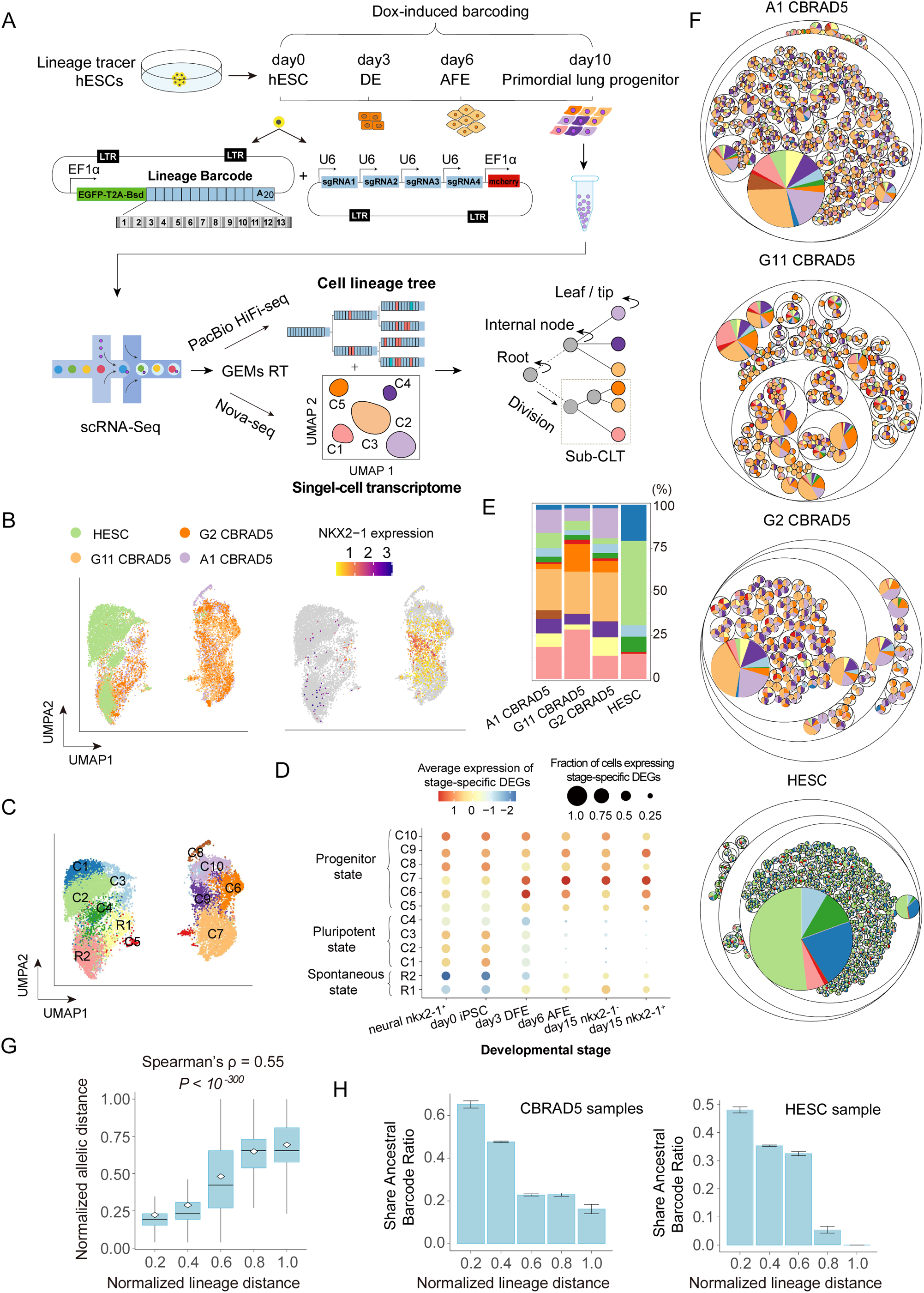
Cell lineage tracing for directed differentiation of primordial lung progenitors. (**A**) Schematic diagram illustrating the overall experimental process. A 10-day directed differentiation from several Lineage Tracer hESCs to primordial lung progenitors (PLP) was conducted along out with simultaneous lineage tracing utilizing inducible CRISPR-Cas9 editing of an expressed lineage barcode (13 editable sites). The resulting colony was assayed for single-cell transcriptomes by Nova-seq and lineage barcode by PacBio HiFi-seq, which were used to reconstruct CLTs with single-cell transcriptomes assigned to tips. (**B**) The variation among single-cell transcriptomes captured in the four samples (one non-differentiating “HESC” sample and three differentiating samples) as shown by UMAP. A data point represents a cell, which is colored based on its source sample on the left panel and the expression level of Nkx2-1 (the marker for PLP) on the right panel. (**C**) Major clusters of the single-cell transcriptomes are differentially colored and labeled by their corresponding cell types. (**D**) In the 12 major cell types (*y* axis), differentially expressed genes (DEGs) found in bulk samples of specific developmental stages preceding PLP (*x* axis) were examined for their average expression levels (dot color) and fraction of cells that expressed the gene (dot size). (**E**) For each of the four samples (x axis), the percentage of cells belonging to each type was shown. The cell types are colored identically to those in panel C. (**F**) Reconstructed CLTs are visualized as circle packing charts for the four samples. Circles represent sub-CLTs, whose sizes indicate the number of terminal cells in the sub-CLTs, while the color (same as panel C) indicates the fraction of terminal cells belonging to each cell type. (**G**) A pair of cells’ normalized lineage distance (the number of internal nodes on the path from one cell to the other, divided by the maximal lineage distance found in the sample) is highly correlated with the normalized allelic distance of their lineage barcodes (the total number of target sites that differed from the reference, divided by the maximum value of 26). All cell pairs were separated into five groups based on their normalized lineage distance (*x* axis), and the distribution of normalized allelic distances (*y* axis) within each group is shown in the form of a standard boxplot, with the mean value indicated by the white point. On top, Spearman’s ρ and *P* value for raw data are indicated. (**H**) The probability of finding a common ancestral allele (as yet-to-decay transcripts) between a pair of single-cell tips decreased as their normalized lineage distance (*x* axis) increased. The error bars indicate the standard error estimated by bootstrapping the cell pairs for 1,000 times.

To assess the CLT of the cultured cells, we employed a modified scGESTALT method^23,31^, which combines inducible cumulative editing of a lineage barcode array by CRISPR-Cas9 with large-scale transcriptional profiling using droplet-based single-cell RNA sequencing. Briefly, we initiated the editing of the lineage barcode concurrently with the directed differentiation using a Cas9 inducible by doxycycline. We used an EGFP-fused cell lineage barcode that consists of 13 editing sites, each of which is targeted by one of four mCherry-fused sgRNAs each containing 2 to 3 mismatches in order to avoid large deletions resulting from excessive editing (**Figure 1A**). These sgRNAs were designed to not target any part of the normal human genome other than the integrated lineage barcode (**Table S1**, see **Methods**). The hESCs carrying the lineage tracing system were subjected to the ten-day directed differentiation, then the colonies were processed for cDNA libraries using the standard 10x Chromium protocol. Each cDNA library was split into two halves, with the first half subjected to conventional RNA-seq for single-cell transcriptomes, and the other half subjected to amplification of the lineage barcode followed by PacBio Sequel-based HiFi sequencing of the lineage barcode (**Figure 1A**).

We obtained single-cell transcriptomes of 3,576/4,400/1,456/5,659 cells respectively from three differentiating colonies CBRAD5-A1/G2/G11 and one parallel non-differentiating hESC colony, all of which appeared to have good quality (**Table S2**). The UMAP clustering of the single-cell transcriptomes revealed a large fraction of cells from differentiating/CBRAD5 colonies separated with those from hESC colonies, clearly indicating their differentiated cell states (**Figure 1B**). We identified 12 major functional clusters within the sampled cells (**Figure 1C**; See **Methods**). These clusters displayed transcriptional states largely compatible with known cell types occurred during the directed differentiation^32^ (**Figure 1D**), and were differentially distributed between hESC and CBRAD5 samples (**Figure 1E**), thereby suggesting successfully induced differentiation and accurate measurement of single-cell transcriptomes. After confirming the sequencing quality of PacBio (**Table S3**), the CLT of each sample was constructed based on the lineage barcode using maximum likelihood method (**Figure 1A/F**; See also **Methods**, **Table S4/S5/S6**). The hierarchical population structures of the colonies were complex and intricate. In support of the accuracy of the CLT, cells more closely related to one another displayed more similar lineage barcode alleles (**Figure 1G**), and are more likely to share yet-to-decay transcripts of ancestral lineage barcode (**Figure 1H**). In conclusion, our experiment reliably captured the coarse-grained phylogenetic relationship of the cells within each colony.

### The cell lineage trees recapitulate key features of the transcriptome divergence

To better elucidate the divergence between the single-cell transcriptomes in the context of the observed clusters, we identified differentially expressed genes (DEGs) in previously published microarray-based transcriptome^33^ data of samples from six timepoints of directed differentiation towards PLP (**Figure 2A**). Note here that despite being sampled on day12, the neural Nkx2-1^+^ transcriptome has been shown to be most similar to that of day 0 hESCs^33^. The Gene Ontology terms enriched with these microarray-based stage-specific DEGs (**Table S7**) were then individually examined for overall activities in our single-cell transcriptomes by the member genes’ average expression levels in each cluster (**Figure 2B**. See **Methods**). For pluripotent stage cells (C1/C2/C3/C4), significantly enhanced activities were found among GO terms enriched with DEGs of day 0/3 samples (including neural Nkx2-1^+^)(**Figure 2B**). The same observations were made for progenitor stage cells C6/C10 in GO terms related to day3/day6 samples, as well as C7/C9 cells in GO terms related to day6/day15 lung samples (**Figure 2B**). These results indicate that the single-cell transcriptomes recapitulated major differentiation stages of the *in vitro* PLP differentiation.

**Figure 2.**
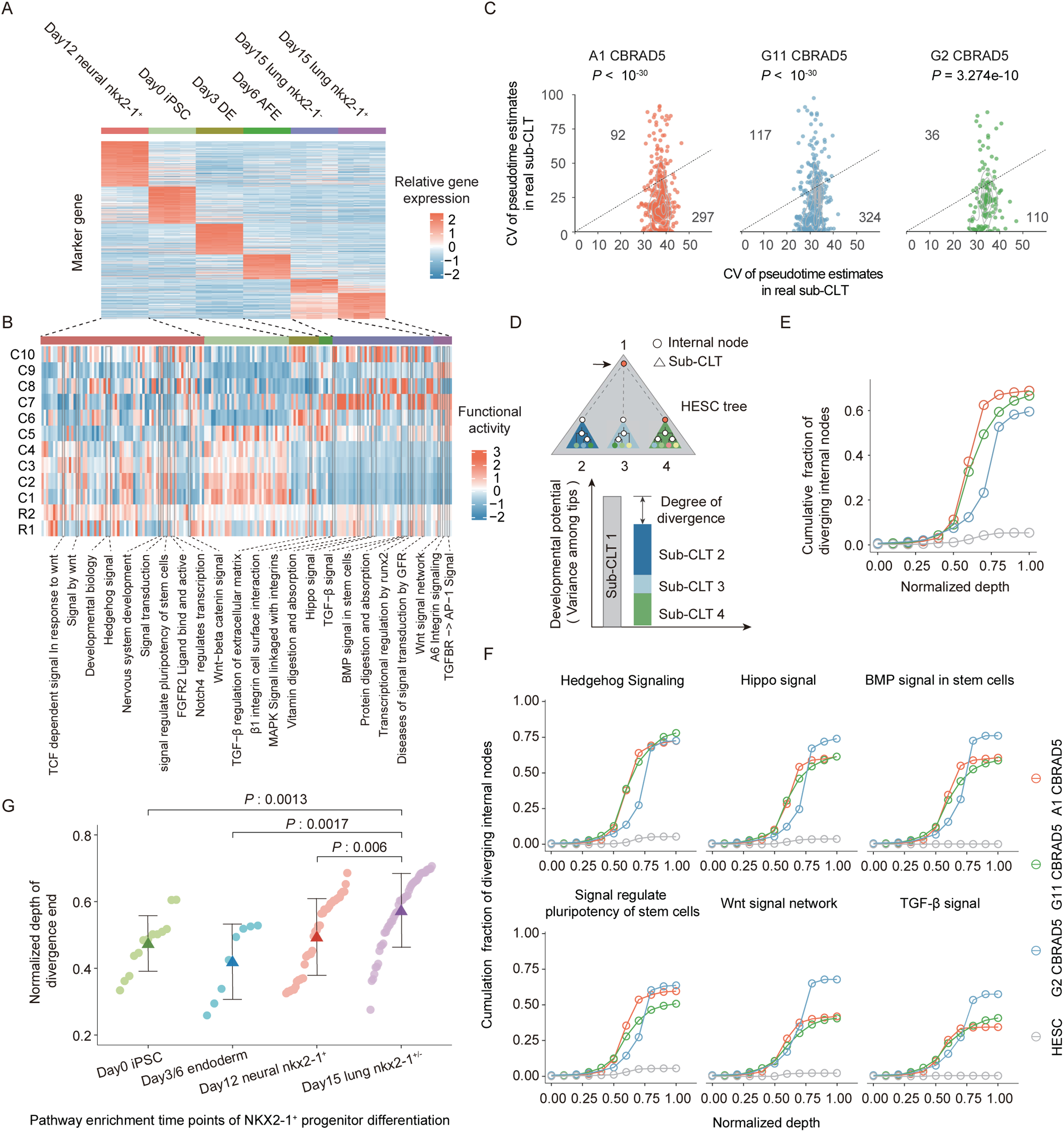
The transcriptome divergence among cell type clusters and among subclones. (**A**) Heatmap for expression levels of DEGs extracted from microarray-based transcriptomes of specific developmental stages (color bars on top) of the directed differentiation^33^. (**B**) Functional activities of GO terms (*x* axis. Full list in **Table S7**) enriched with stage-specific DEGs were shown for every cluster (*y* axis) identified in our samples. Here functional activity as indicated by the color scale was estimated by the average Z-score-transformed expression of all genes annotated with the GO term. (**C**) A coefficient of variation (CV) was calculated using pseudotime estimates of single-cell transcriptomes within a sub-CLT. These CVs were plotted for all real sub-CLTs (y axis) and corresponding randomized sub-CLTs generated by shuffling all tips (x axis) in each differentiating sample (name on top). As the dashed line indicates *x* = *y*, sub-CLTs with CVs lower than random expectation (i.e. restricted variation) will appear below it. Each panel includes the number of CLTs above and below the dashed line, which was also tested against the binomial expectation (50% below the line) and yielded the *P* values on top. (**D**) Schematic diagram for the PERMANOVA-based estimation of transcriptome divergence for an internal node. (See **Methods**) (**E**) Cumulative fraction (*y* axis) of internal nodes exhibiting significant transcriptome divergence as the normalized depth (*x* axis) considered increased. Results from different samples were shown with different colors, as indicated by the color legend. Dashed line indicates *y* = 0.05. (**F**) Same as panel **E** except that the analyses were limited to specific GO terms indicated on top of each panel. (**G**) The normalize depths at which the divergence of specific functions initiated. GO terms enriched of day3/day6/day15 cell type marker genes were examined (*x* axis), with the exception of five terms whose divergence signal was not higher in differentiating versus non-differentiating samples. The GO terms listed in panel **F** were indicated by a black arrow.

Our data also permit us to resolve divergence among sub-CLTs. It is commonly understood that the developmental process involves an increase in transcriptional divergence among cells and a reduction of developmental potentials in individual cells. Analyzing single-cell transcriptomes among sub-CLTs should reveal these patterns with fine resolution, especially when using high-density CLTs as we obtained. As an initial assessment for whether there is transcriptional divergence among sub-CLTs in the differentiating samples, we calculated for each sub-CLT, the CV (coefficient of variation) of the pseudotime estimates^34^ (see **Methods**) of all its tips. When compared with their null expectations generated by randomly shuffling all tips, majority of these CVs were significantly smaller (**Figure 2C**), suggesting cells in the same sub-CLT are more similar than expected by the full range of transcriptional variation, an observation directly supports the transcriptional divergence among sub-CLTs.

For a more detailed analyses, we quantified the developmental potential of an internal node by the multivariate variance among its descendant single-cell transcriptomes, which then allowed us to perform PERMANOVA-based statistical tests (PERmutational Multivariate Analysis Of VAriance, see **Methods**) for the transcriptomic divergence. Briefly, by subtracting from the developmental potential of a focal node by the sum of the potentials of all its daughter nodes, we estimated the degree of divergence that occurred during the growth of the focal node (**Figure 2D**). Using the degree of divergence seen in the hESC sample as the null distribution, an average of ∼65.1% internal nodes of the CBRAD5 samples displayed significant divergence (**Figure 2E**), whereas only ∼5.4% internal nodes displayed divergence in the HESC sample. When such degree of divergence is depicted against normalized depths (**Methods**) of the corresponding nodes, the CBRAD5 samples consistently showed rapid divergence that is not seen in HESC samples (**Figure 2E**). Please note that divergence here is not equivalent to differentiation, since two sister cells differentiating into the same fate would not reveal any divergence for their mother cell. In other words, divergence implies asymmetric division creating daughter cells of different developmental potentials, whereas differentiation can occur during symmetric division giving rise to a pair daughter cells that both activate a particular function or differentiate in the same direction.

By restricting the above analysis to gene subsets associated with specific GO terms, it is possible to elucidate the progression of divergence in the corresponding cellular functions. As shown in several key GO terms including Wnt signaling, the cumulative growth in the fraction of internal nodes with significant divergence at various normalized depths is also highly reproducible among CBRAD5 samples, and it differs from the hESC sample (**Figures 2F**). Additionally, we examined whether our CLT data could resolve the temporal order of divergence completion for different cellular functions. To this end, we traced all root-to-tip paths on the CLTs and calculated the average depth of the last (furthest from the root) internal node exhibiting significant divergence on a pathway. As a result, the normalized depths of divergence completion appear consistent with known temporal orders of key developmental events (**Figure 2G**). Collectively, these results indicate that our dataset of single-cell transcriptomes and CLTs allowed the elucidation of cellular development with reasonable resolution.

### Transcriptional memory has limited contribution to developmental canalization

Following confirmation of the CLT data’s resolution, we began searching for contributors to developmental robustness using CLTs. A first hypothesis is that transcriptional memory may have constrained gene expression variation during development, which would canalize transcriptomic state during development and contribute to robustness. In this context, transcriptional memory is the phenomenon of cells closely related on the CLT displaying similar expression levels due to the inheritance of the same cellular contents (proteins/transcripts) and/or epigenetic states from recent common ancestors^29,31,35,36^. Nevertheless, gene expression can also be restricted by transcriptional regulation that has nothing to do with cellular inheritance, such as negative feedback^37^ and denoising promoters^38^. If the transcriptional memory dominates the experimented differentiation, one would expect all cells of the same type would have been clustered into an exclusive sub-CLT, which is clearly not the case (**Figure 1F**). For a quantitative analyses, we reasoned that the CV of single-cell expression levels within real sub-CLTs should reflect the combined effect of transcriptional memory and inheritance-independent regulation (**Figure 3A** top), whereas that of CLTs randomized by shuffling cells of the same type at different lineage positions should reflect only inheritance-independent regulation but not transcriptional memory (**Figure 3A** bottom). It is therefore possible to isolate the contribution of transcriptional memory to the expression constraint by contrasting the CV of real CLTs with that of randomized CLTs (**Figure 3A** and **Methods**), which is hereinafter referred to as the “memory index”. We note that this definition of memory index is similar to that used in previous transcriptional memory-related studies ^29,39^.

**Figure 3.**
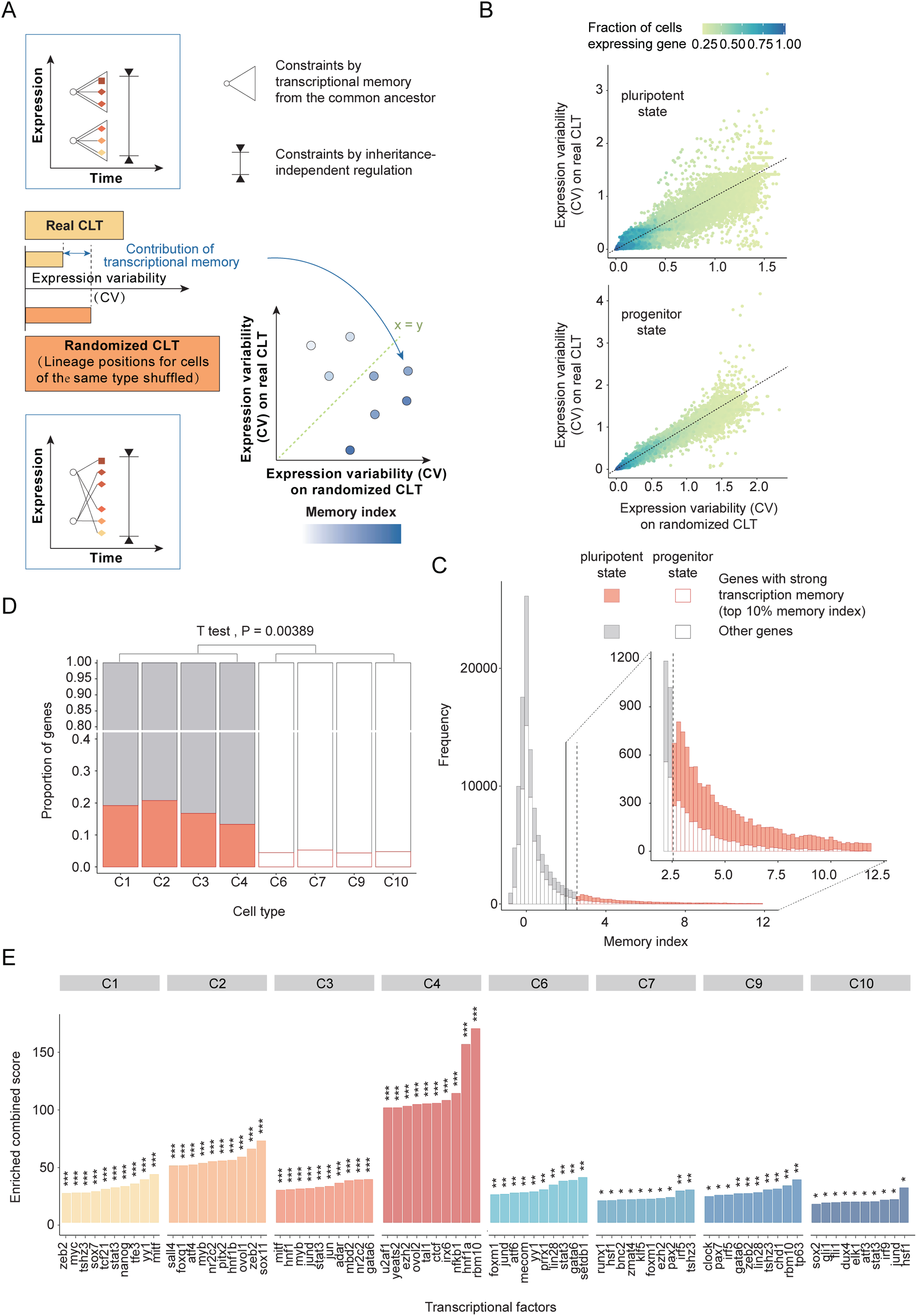
Limited contribution of transcriptional memory in differentiation. **(A)** Schematic diagram for the CLT-based estimation of transcriptional memory. **(B)** Expression variability in the real CLT (*y* axis) compared to that in the randomized CLT (*y* axis). Each dot represents a gene in a cell type. Dot color shows the fraction of cells within the cell type that express the gene, as indicated by the color scale on top. (**C**) A stacked histogram showing the distribution of the memory indices calculated. A filled bar represents those estimated from pluripotent cell types and an empty bar represents those estimated from progenitor cell types. Genes exhibiting strong transcriptional memory, i.e. those with a memory index ranking among the top 10%, were red, while others were gray. The inset shows a zoomed-in view of the large memory index region. (**D**) Among different cell types, the fraction (height of bar) of genes exhibiting high memory indices was compared. The bars are colored similarly to those in panel C. (**E**) Gene sets responsive to perturbation of individual transcription factors (*x* axis) were tested for the enrichment of genes exhibiting strong signal of transcriptional memory (see **Methods**). The top ten transcription factors with the highest combined enrichment score (*y* axis) were shown for each cell type. The statistical significance of enrichment according to Fisher’s exact test is indicated as *:*P*<0.05; **:*P*<0.01; ***:*P*<0.001.

For each cell type, we calculated an overall memory index for each gene in each sub-CLT (**Figure 3B**). The top (10%) memory indices (**Figure 3C**) were found to be enriched in pluripotent cell types (C1/C2/C3/C4) as compared to progenitor cell types (C6/C7/C9/C10) (*t*-test *P*=0.0039, **Figure 3D**), suggesting that transcriptional memory is more important to maintaining pluripotency than differentiation. Because transcriptional memory is mediated by cellular contents inherited from mother to daughter cells, such as transcription factors, we hypothesized that these genes with top memory indices should be enriched downstream of some transcription factors. Thus, we tested these genes for enrichment in genes responsive to genetic perturbation of individual transcription factors^40^ (see **Methods**), and made two observations. First, some transcription factors with known involvement in the experimented differentiation, such as Nanog in the pluripotent C1^41^ and Gata6 in progenitor C6/C9^42^, indeed exhibit significant enrichment of the genes with top memory index. Second, the enrichment was generally stronger for pluripotent cell types than it was for progenitor cell types (**Figure 3E**), a pattern again suggesting that transcriptional memory only played a minor role in differentiation, which is at least not as significant as in maintaining pluripotency.

### Stable cell type compositions across sub-clones supports robust development

Observations above indicate that terminal cells within a sub-CLT have restricted fates that are not dominated by transcriptional memory from the common ancestor (root of the sub-CLT). This observation automatically prompted an assessment of the cell fate restriction imposed by inheritance-independent regulation, as well as its contribution to the robustness of developmental processes. We reasoned that inheritance-independent regulation should result in multiple similarly restricted sub-CLTs dispersed across the entire CLT. Therefore, we calculated the terminal cell type composition for each sub-CLT found in the CBRAD5 samples and compared it with the overall composition of the corresponding full CLT (see **Methods**). Intriguingly, the cell type compositions of sub-CLTs are usually more similar to those of the full CLTs than expected in randomized CLTs (**Figure 4A-C**). A closer examination of some sub-CLTs reveals a highly stable terminal cell type compositions. For example, there are 35 sub-CLTs that generated subclones with highly stable (<10% deviation) proportions of 0.13, 0.39, 0.13 and 0.18 respectively for C6, C7, C9 and C10 (the top four progenitor cell types), which corresponds to the average proportion of these cell types in the three differentiating samples (**Figure 4D**). This observation suggests that a stereotyped developmental program may exist that produces subclones with highly similar compositions of cell types derived from multiple ancestral cells.

**Figure 4.**
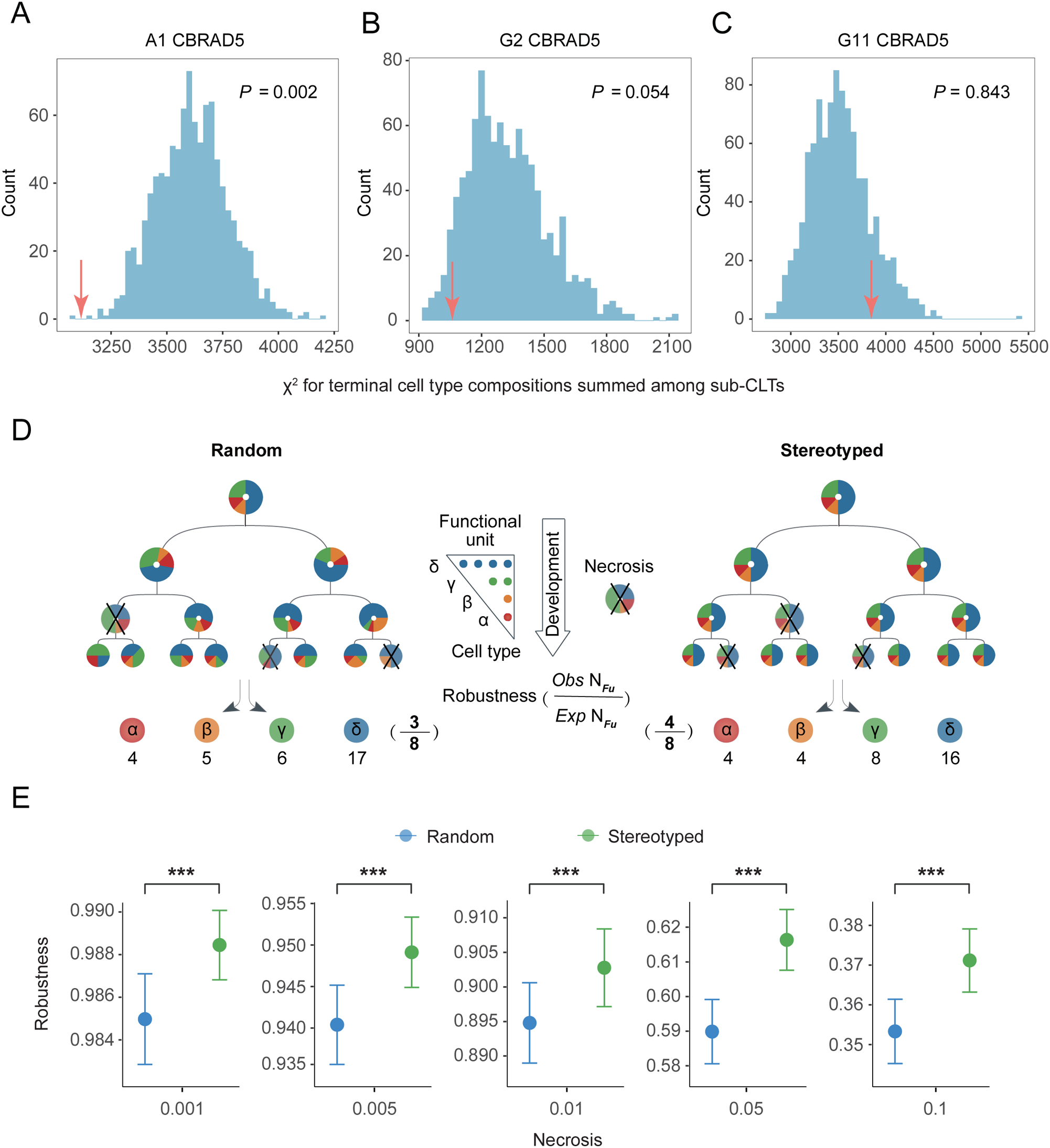
Stable cell type composition across sub-clones supports robust development. **(A-C)** In each panel for each of the CBRAD5 samples (names on top of the panel), the diversity of compositions of terminal cell types within sub-CLTs were estimated by a summed chi-square value (*X*^2^) (see **Methods**) as indicated by the red arrows. The same summed *X*^2^ values were calculated for 1,000 randomized CLTs, whose distribution was shown as a blue histogram. The probability of a summed *X*^2^ value being smaller than the observation (red arrow) is indicated by the *P* values in the panel. **(D)** A schematic diagram showing a simple model of the functional robustness of the random (left) versus stereotyped (right) development against random necrosis (indicated by “X”). The robustness is quantified by the number of functional units (with cell type compositions indicated in the triangle) that can be formed by terminal cells surviving necrosis, as exemplified at the bottom. **(E)** Robustness (*y* axis) of the random (blue) *versus* stereotyped (orange) development under different rate of necrosis (*x* axis), as estimated by the model in **D**.

The observed polyclonal stereotypic development can be understood from two perspectives. On the one hand, the consistent execution of such a developmental program across subclones may be by itself a manifestation of robust genetic and/or molecular regulation. On the other hand, stable cell type compositions across subclones might enhance developmental robustness. We examined this latter perspective by simulating a CLT for the development of a single cell into an “organoid” consisting of 1,024 cells (i.e., 10 cell cycles) comprised of four types (namely α, β, γ, and δ) of cells in a 1:1:2:4 ratio. These cells formed 128 functional units each consisting of one α cell, one β cell, two γ cells, and four δ cells. Normally developed organoid consisting of 128 functional units (assuming sufficient cellular migration) are considered 100% functional. Meanwhile, CLT perturbed by random necrosis (see below), which results in the loss of some ancestral cells and all their descendants, has a functional capacity defined as the fractional survival rate of functional units with proper cellular composition. This design was inspired by the observation that functional units in living tissues, such as mouse pancreatic islets, display a highly stable cell type composition as the outcome of normal development^26^. To generate the normal (necrosis-free) CLT with the predetermined number of cells of each type, three models were used. The first “random” model assigns each cell to a random tip of the CLT regardless of its cell type (**Figure 4D** left). A second “stereotyped” model defines all eight-tip sub-CLTs as strictly consisting of one α cell, one β cell, two γ cells, and four δ cells, but different placements of these cells are allowed on the tips (**Figure 4D** right). A total of 1,000 normal CLTs were generated under each model, and the functional capacity of each CLT was determined by exposing all (internal or terminal) cells to 2% random necrosis. As compared to the random model, we found that CLTs generated with the stereotyped models were always more resilient to necrosis (**Figure 4E**). Such enhanced developmental robustness is more evident at higher rate of necrosis (**Figure 4E**). This simulation suggests that the observed stable cell type composition among subclones is associated with increased developmental robustness.

### Stereotyped cell lineage trees underlie stable cell type compositions

We next seek further evidence for the existence of stereotyped developmental programs based on the CLT data at hand. Specifically, we hypothesized the existence of multiple sub-CLTs with highly similar topology and terminal cell types. As recurrent sub-sequences of biological sequences, such as transcription factor binding sites, are usually referred to as “sequence motifs”, we call our target recurrent sub-CLTs “tree motifs” or simply “motifs”.

Just as sequence motifs are identified by comparisons between (sub-)sequences, tree motifs should also be identified through comparisons between (sub-)CLTs. In order to identify potential tree motifs in the CLT of the differentiating samples, we utilized Developmental cEll Lineage Tree Alignment (DELTA), an algorithm we previously developed for quantitative comparisons and alignments between CLTs^43^ (**Figure 5A**, see **Methods**). Using a dynamic programming scheme analogous to that employed by classical algorithms looking for similarities between biological sequences (e.g. the Smith-Waterman algorithm), the DELTA algorithm searches for pairs of homeomorphic sub-CLTs^43^ within two given full CLTs. It is important to emphasize that a high-density CLT is essential for DELTA analyses because it would be impossible to identify any motif with only a very small fraction of cells being sampled in each sub-CLT.

**Figure 5.**
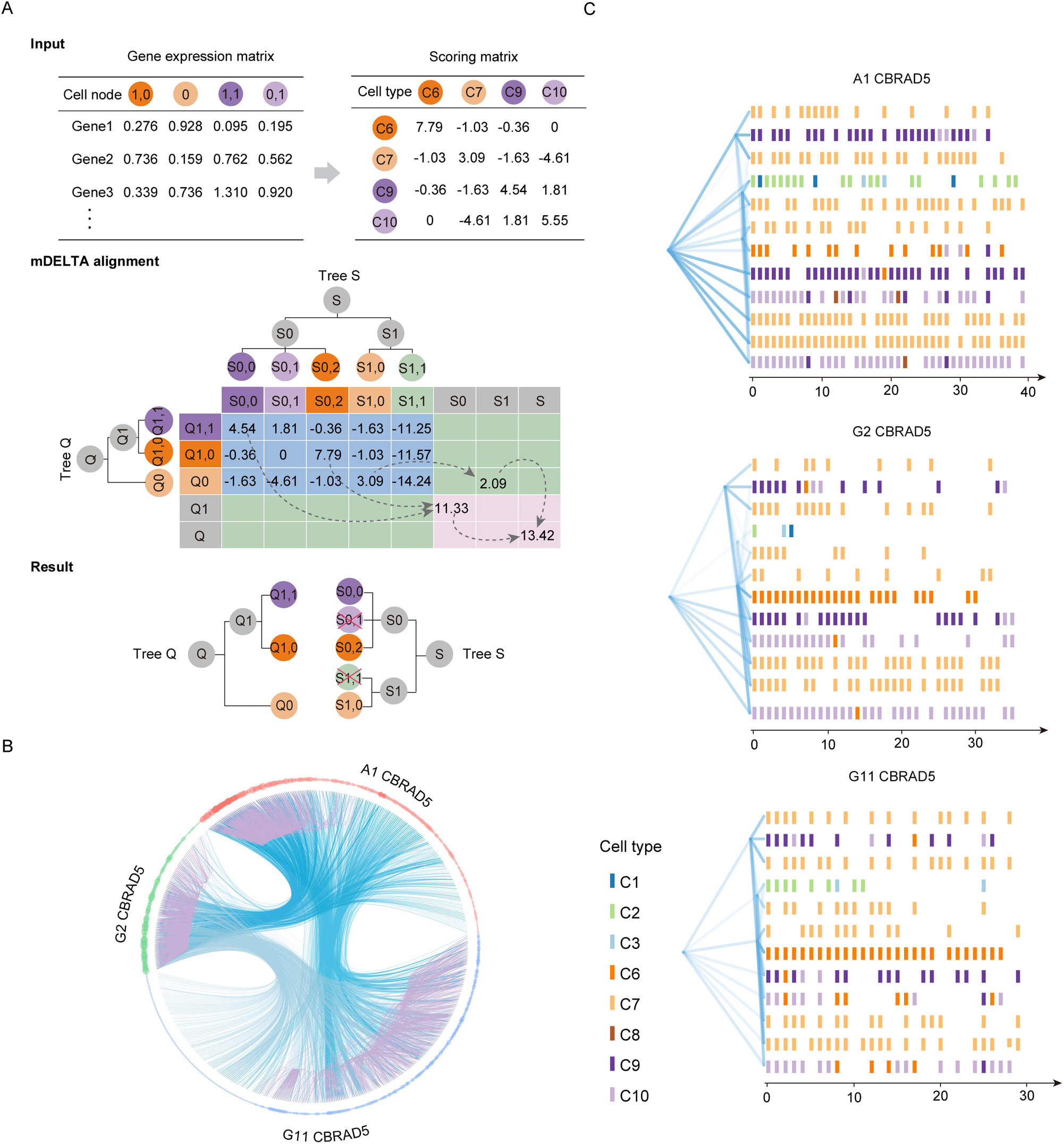
The heritability and of the stereotyped developmental program. **(A)** The input (left) for DELTA includes two CLTs and the expression profiles of all terminal cells on these CLTs. DELTA uses a dynamic programming procedure (middle) to compare the two CLTs and identify homeomorphic sub-CLTs. The procedure has three phases, including (i) a cell pair scoring stage, (ii) a forward stage that maximizes the alignment scores by finding the best correspondence between terminal cells, and (iii) a backtracking stage for extracting the alignment behind the maximized scores. The output (right) is one or more aligned sub-CLTs ordered by decreasing alignment scores. See **Methods** for more details. (**B**) A circular plot of the top 1,000 sub-CLT pairs found by DELTA in each of the six pairwise comparisons among the CLTs from the three differentiating samples. In the outer circle, each sub-CLT is represented by a dot, with the color indicating its source sample. Each pair of homeomorphic sub-CLTs identified by DELTA is shown by curved links between two corresponding dots, where inter-sample pairs are represented by blue links and intra-sample pairs by purple links, respectively. A dot’s size indicates how many links it has. Only sub-CLTs with at least one link are included. (**C**) One highly recurrent tree motif found in all three samples is shown by “densitree” plots. All sub-CLTs homeomorphic to a specific reference sub-CLT are extracted from DELTA results in panel **B**. They were separated by their source sample as indicated on top of each plot. In each plot, the DELTA-aligned topological structure of each sub-CLT (including the reference sub-CLT) is drawn with transparency on the left so that common topologies can be seen as darker lines. Each column of tiles on the right shows the DELTA-aligned terminal cell types on one of the homeomorphic sub-CLTs, by the colors listed to the left of the plot. The left-most column of tiles is always the reference sub-CLT. The number at the bottom indicates the number of sub-CLTs found as homeomorphic to the reference sub-CLT.

Consistent with our prediction, DELTA identified a large number of highly similar sub-CLT pairs between and within differentiating samples (**Figure 5B**). Some of the most frequently occurring sub-CLTs exhibited a consistent structure, comprising multiple layers of internal cells and a stable composition of terminal cell types (**Figure 5C**). Groups of such highly similar sub-CLTs represent strong candidates of tree motifs on the developmental CLT, and strongly supports the existence of a stereotyped developmental program that contributes to developmental robustness.

## Discussion

In the current study, we have reconstructed high density developmental CLTs for *in vitro* directed differentiation from hESC to primordial lung progenitors. In comparison with CLTs of non-differentiating hESC colonies, differentiation CLTs showed a clear signal of transcriptomic divergence that recapitulates known involvements of key developmental regulatory pathways. Using CLTs, we investigated mechanisms that might have contributed to developmental robustness at the intercellular level. It was found that transcriptional memory had limited effects on canalizing cell fates within subclones. Nevertheless, we found that multiple subclones exhibit stable compositions of terminal cell types, which enables sufficient numbers of cells in proper composition to be generated, and thus, a more robust development. By using a CLT alignment algorithm, we further showed that the observed stable cell type composition is underlied by stereotyped sub-CLTs with similar topology and terminal cell fate. Collectively, our results show that stereotyped sub-CLTs support robust development.

Beyond the specific mechanisms underlying developmental robustness, our findings suggest a novel perspective regarding cell types within the context of stereotyped sub-CLTs. In particular, just as letters can be better understood within the context of words, and nucleotides/amino acids can be better understood within the context of sequence motifs, stereotyped sub-CLTs can potentially bridge our knowledge of the atlas of cell types and their organization into functional tissues.

## Methods

### Design of the lineage tracer hESC cell line

To design the lineage barcode and corresponding sgRNA, we first generated randomized 20-bp candidate sgRNA sequences with >3 substitutions relative to any human genome fragments. Among these candidates, the spacer sequence 5’-TATTCGCGACGGTTCGT-ACG-3’ was selected as sgRNA1. A total of 13 protospacer sequences were designed based on sgRNA1 according to the following criteria: (i) each protospacer contained 2-3 mismatches with sgRNA1, (ii) there was no recurrence of any sequence of 9 bp or longer, and (iii) consecutive repeats of the same nucleotide for more than 2 bp were completely absent. The 13 protospacers (along with PAM, or protospacer adjacent motif) were organized according to decreasing CFD (cutting frequency determination) scores into the full lineage barcode ^44,45^. The next three sgRNAs, sgRNA2, sgRNA3, and sgRNA4, were designed to perfectly match the 9th, 12th, and 13th protospacers, but with lower CFD scores (<0.55) for other protospacers, because these three protospacers were rarely edited in preliminary experiments using only sgRNA1. To facilitate capture by poly-dT reverse transcription primers on 10x gel beads, the full lineage barcode with a 20-nt poly-dA(A20) 3’ tail was inserted into the 3’UTR of an EGFP driven by an EF1α promoter.

We constructed lineage tracer hESC cell lines by genomic integration of the lineage barcode, doxycycline-inducible Tet-on Cas9 and the sgRNAs. Briefly, the lineage barcode vector (pLV-EF1A>EGFP:T2A:Bsd:V1, VectorBuilder, no:VB1709 11-1008qmt) was constructed by the Gateway system and then transfected into H9 hESCs with MOI=0.15. The EGFP-fused lineage barcode was confirmed to exist as a single copy in the genome and to be highly expressed after blasticidin selection (15 g/ml, InvivoGen, no. ant-bl-1) and flow cytometry sorting. Then the Tet-on inducible Cas9 vector (PB-Tet-ON-T8>Cas9:T2A:puro-PGK:rtTA, donated by Professor Jichang Wang, Zhongshan School of Medicine, Sun Yat-sen University) was co-transfected with hyPBase (VectorBuilder, no: VB190515-1005nrp) in a ratio of 100ng:1ug for 1×10^7^/ml cells by Neon^TM^ transfection system (Life, MPK5000). In order to ensure adequate Cas9 expression for efficient editing, we applied double reinforced selection of Puromycin (1.0 μg/ml, InvivoGen, no. ant-pr-1) and Doxycycline (Dox, 1.0μg/ml, sigma, D9891-1G) for 7 days. Lastly, the sgRNA vector (pLV-U6>sgRNA1>U6>sgRNA2>U6>sgRNA3>U6>sgRNA4-EF1α>Mcherry:T2A:Neo,VB1912 11-3149jwe) was constructed by Golden Gate ligation and transfected at MOI=30. H9 hESC cells with high expression of sgRNAs (fused with mCherry) were enriched by G418 selection (1000 μg/ml, InvivoGen, ant-gn-1) for 11 days and flow cytometry sorting. Expression levels of Cas9, lineage barcode and sgRNA1 transcripts were detected by RT-qPCR with primers listed in **Table S8**.

The editing efficiency of the lineage tracer hESC cell line was evaluated by inducing Cas9 expression in mTesR media with 1.0 µl/ml Dox for five days. We extracted gDNA from all cells using DNeasy Blood & Tissue Kits (Qiagen, no.69504). Using primers gDNA-V1f and gDNA-V1r (**Table S8**), we amplified the lineage barcode from gDNA using Phanta Max Super-Fidelity DNA Polymerase (Vazyme, No. P505), which was then cloned into pCE-Zero (Vazyme, No. C115)., which was then cloned into the pCE-Zero vector (Vazyme, No. C115). The efficiency of editing was then evaluated by colony PCR and Sanger sequencing for 50 recombinant clones.

Additionally, we examined editing efficiency in the context of our directed differentiation experiment, in which only a small number of initial cells were used to form each colony. In four 96-well dishes, matrigel (Corning, No. 354277) was plated and each well was seeded with < 10 log-phased lineage tracer hESC cells manually by micromanipulation. For 11 days, the cells were cultured in 100 µl of mTesR media, to which 10 µl of cloneR (Stemcell, No.05888) were added on day0 and day2, and 1.0 µg/ml Dox+ mTesR media was added and refreshed every 48 hours since day2. Normally surviving colonies after the 11-day culture were harvested by GCDR (Stemcell, No.07174). Next, 50ng of genomic DNA was extracted from each colony using the QIAamp DNA Micro Kit (Qiagen, No.56304) and PCR amplified for the lineage barcode. The Cas9-induced mutations accumulated during colony formation were then identified by Sanger sequencing, TA cloning or Illumina HiSeq PE250 sequencing. Specifically, the raw HiSeq data were trimmed by fqtrim (https://ccb.jhu.edu/software/fqtrim/) with default parameters. The paired reads were merged by FLASH^46^ using 30 bp of overlapping sequence and 2% mismatches. Sequences alignable to the human reference genome by Bowtie2 with default parameters^47^, or to primer sequences of gDNA-V1f and gDNA-V1r with two mismatches, were removed as they likely represented nonspecifically amplified sequences. MUSCLE^48^ aligned the sequenced lineage barcode to the wild-type lineage barcode using default parameters. The editing events of each sequence were identified according to a previous method^44^.

### Validating directed differentiation from hESC to lung progenitor and alveolosphere

Using the BU3 NGST (NKX2-1GFP; SFTPCtdTomato) iPS cell line (donated by Professor Darrell N. Kotton, Deparment of Medicine, Boston University), we tested the protocol of directed differentiation towards lung progenitor and alveolosphere published by Kotton and colleagues^30^. Briefly, in six-well dishes pre-coated with Matrigel (Stemcell, No.356230), 2×10^6^ cells maintained in mTESR1 media were differentiated into definitive endoderm using the STEMdiff Definitive Endoderm Kit (StemCell, No.05110), adding supplements A and B on day 0, and supplements B only on day 1 to day 3. Flow cytometry was used to evaluate the efficiency of differentiation to definitive endoderm at day 3 using the endoderm markers CXCR4 and c-KIT (Anti-human CXCR4 PE conjugate, Thermo Fisher, MHCXCR404,1:20; Anti-human c-kit APC conjugate, Thermo Fisher, CD11705, 1:20;dilution. PE Mouse IgG2a isotype, Thermo Fisher, MG2A04,1:20; APC Mouse IgG1 isotype, Thermo Fisher, MG105, 1:20) based on the method of Sahabian and Olmer^49^. After the endoderm-induction stage, cells were dissociated for 1-2 minutes at room temperature with GCDR and passaged at a ratio between 1:2 to 1:6 into 6 well plates pre-coated with growth factor reduced matrigel (Stemcell, No.356230) in ‘‘DS/SB’’ anteriorization media, which consists of complete serum-free differentiation medium (cSFDM) base, including IMDM (Thermo Fisher, No.12440053) and Ham’s F12 (Corning, No. 10-080-CV) with B27 Supplement with retinoic acid (Gibco, No.17504044), N2 Supplement (Gibco, No.17502048), 0.1% bovine serum albumin Fraction V (Sigma, A1933-5G), monothioglycerol (Sigma, No. M6145), Glutamax (ThermoFisher, No. 35050-061), ascorbic acid (Sigma,A4544), and primocin with supplements of 10 μm SB431542 (‘‘SB’’; Tocris, No.1614) and 2 μm Dorsomorphin (‘‘DS’’; Sigma, No. P5499). In the first 24 hours following passage, 10 μmY-27632 was added to the media. After anteriorization in DS/SB media for three days (72 hr, from day 3 to day 6), cells were cultured in “CBRa” lung progenitor-induction media for nine days (from day 6 to day 15). This CBRa media consists of cSFDM containing 3 μm CHIR99021 (Tocris, No.4423), 10 ng/mL rhBMP4 (R&D, 314-BP-010), and 100 nM retinoic acid (RA, Sigma, No. R2625). At day 15 of differentiation, single-cell suspensions were prepared by incubating the cells at 37°C in 0.05% trypsin-EDTA (Gibco, 25200056) for 7-15 minutes. The cells were then washed in media containing 10% fetal bovine serum (FBS, ThermoFisher), centrifuged at 300 g for 5 minutes, and resuspended in sort buffer containing Hank’s Balanced Salt Solution (ThermoFisher), 2% FBS, and 10 μm Y-27632. The efficiency of differentiation into NKX2-1^+^ lung progenitors was evaluated either by flow cytometry for NKX2-1-GFP reporter expression, or expression of surrogate cell surface markers CD47^hi^/CD26^lo^. Cells were subsequently stained with CD47-PerCPCy5.5 and CD26-PE antibodies (Anti-human CD47 PerCP/Cy5.5 conjugate, Biolegend, Cat#323110, 1:200; Anti-human CD26 PE conjugate, Biolegend, Cat#302705, 1:200; PE mouse IgG1 isotype, Biolegend, Cat#400113, 1:200, PerCP/Cy5-5 mouse IgG1 isotype, Biolegend, Cat#400149, 1:200) for 30 min at 4 °C, washed with PBS, and resuspended in sort buffer based on the method of Hawkins and Kotton ^49^. Cells were filtered through a 40 μm strainer (Falcon) prior to sorting. The CD47^hi^/CD26^lo^ cell population was sorted on a high-speed cell sorter (MoFlo Astrios EQs) and resuspended in undiluted growth factor-reduced 3D matrigel (Corning) at a dilution of 25-100 cells/μl, with droplets ranging in size from 20 μL in 96 well plates. Cells in 3D matrigel suspension were incubated at 37 °C for 20-30 min, followed by the addition of warm media. The differentiation into distal/alveolar cells after day 15 was performed in ‘‘CK+DCI’’ medium, consisting of cSFDM base, with 3 μm CHIR (Tocris, No.4423), 10 ng/mL rhKGF(R&D, No.251-KG-010) (CK), and 50 nM dexamethasone(Sigma, No. D4902), 0.1 mM 8-Bromoadenosine 3’,5’-cyclic monophosphate sodium salt (Sigma, No.B7880) and 0.1 mM 3-Isobutyl-1-methylxanthine (IBMX; Sigma, No.I5879) (DCI). Immediately after replating cells on day 15, 10 μm Y-27632 was added to the medium for 24 hours. Upon replating on day 15, alveolospheres developed in 3D Matrigel culture outgrowth within 3-7 days, and were maintained in CK+DCI media for weeks. These spheres were analyzed by Z stack live images of alveolospheres taken and processed on the Leica DMi8 fluorescence microscope.

### Directed differentiation followed by simultaneous assessment of single-cell transcriptomes and cell lineage tree

Based on the results from the full directed differentiation experiment above, we aimed to evaluate single-cell transcriptomes and CLTs simultaneously for directed differentiation from hESCs to PLP, a stage at which the colony has <10,000 cells, allowing us to sample a large proportion of cells. To prepare suitable ancestor hESCs, the cell colonies outgrowth after 5-7 days, plated in 96-well dishes with microscopic selection for GFP^+^ mCherry^+^, were digested with GCDR to form ∼50 μm aggregates, and cultured in mTesR media until day 5. Combining selection and induction by dox and puro from day 5 to day 7, the normally survived GFP^+^ mCherry^+^ colonies were capable of Cas9 expression and marked by primary editing events (to distinguish ancestor cells), as confirmed by DNA extraction and barcode PCR and sanger sequencing. The cell colonies with primary editing events were digested by GCDR for cell counting (∼ 4000 cells) and resuspended at a density of 10 cells/μl. 1μl cell suspension was added into each well of 96-well dishes plated with 1:10 diluted Matrigel (Corning, No.354277) for culture in mTesR media with ClonR (10:1) (Stemcell, No.05888) added in the first 48h to promote the survival of very few stem cell. Directed differentiation was then initiated by applying both dox (1.0μg/ml, for editing the lineage barcode) and the STEMdiff Definitive Endoderm Kit to the normally survived colonies. Later stages of directed differentiation followed the differentiation protocols described above, with the exception that it was stopped on the tenth day after its initiation. Finally, colonies with intermediate size (∼ 5,000 cells as approximated by colony size) and ≥50% GFP^+^ Mcherry^+^ cells were digested with 0.05% trypsin-EDTA for 1 minute at 37 °C, washed in PBS containing 10% fetal bovine serum (FBS, ThermoFisher), centrifuged at 500 g for five minutes, and resuspended in single cell resuspension buffer containing PBS and 0.04% BSA. Using the standard 10x Chromium protocol, cDNA libraries were prepared from these single cell suspensions. Each cDNA library was split into two halves, with the first half subjected to conventional RNA-seq for single-cell transcriptomes, and the other half subjected to amplification of the lineage barcode followed by PacBio Sequel-based HiFi sequencing of the lineage barcode (**Figure 1A**).

### Analysis of scRNA-seq

Following the 10x Genomics official guidelines, we used the Cell Ranger^50^ pipeline to map raw reads to the human reference genome (GRCh38) by STAR^51^ and obtained the read counts for each gene. Using Seurat v4.4^52^, we retained cells with <10% mitochondrial reads and >200 expressing unique features detected. Then highly variable genes were detected by Single-cell Orientation Tracing (SOT)^53^, which were then subjected to Principle Component Analysis, followed by batch effect correction by Harmony^54^. We then clustered cells based on the cell-cell distance calculated by FindNeighbors and FindClusters using the Harmony-normalized matrix of gene expression. Then, we used runUMAP for visualization and FindAllMarkers to obtain differentially expressed genes (DEGs) among clusters. To identify cell types, we downloaded microarray data from Gene Expression Omnibus (GEO)^33,55^, and extracted DEGs (Wilcoxon Rank Sum test, *P* < 0.01) in different stages of differentiation towards PLP. We scored the clusters base on the average expression and numbers of expressed stage-specific DEGs. Finally, we named 12 cell cluster based on the inferred order of appearance in the differentiation progress.

### Construction of cell lineage trees

Based on the PacBio HiFi sequencing results, we built and assessed the quality of the CLT from PacBio HiFi reads following our previous pipeline^31^. Briefly, using HiFi-seq raw sequences, we called consensus sequences separately from positive and negative strand subreads from each zero-mode waveguide (ZMW). We reserve only consensus sequences with at least three subreads and identifiable barcode primers (**Table S8**, allowing up to two mismatches). From the consensus sequences, 10x cell barcodes and UMIs were extracted and matched to those from scRNA-seq, with one mismatch allowed. Lineage barcode sequences were then extracted from the consensus sequences, grouped by identical cell barcode and UMI, then merged by MUSCLE alignment followed by selecting the nucleotide with the highest frequency at each site. After MUSCLE alignment of the merged sequence to the reference lineage barcode, the editing events were called^44^. Then, for each lineage barcode allele from the same cell, the frequency was calculated as the total number of UMIs of the allele and its ancestral allele. Here, the ancestral allele of a specific allele was defined as any allele in which the observed editing events were a subset of the editing events in the focal allele. Finally, the lineage barcode allele of a cell was defined as the allele with the highest frequency, prioritizing the alleles with more editing events if the frequencies were equal. For each sample, all cells with a lineage barcode and a single-cell transcriptome were used to construct a multifurcating lineage tree based on the lineage barcode using the maximum likelihood (ML) method implemented by the IQ-TREE LG model ^56^.

### Transcriptome divergence among cell type clusters

To elucidate the transcriptomic divergence among the observed clusters in the context of the directed differention towards PLP, we extracted stage-specific DEGs with the top 10% most extreme fold-change relative to other stages (**Figure 2A**, using microarray data^33^ mentioned above), and identified the Gene Ontology terms enriched (BH-adjusted *P* < 0.01, Fisher’s exact test) with these stage-specific DEGs. After eliminating GO terms that have very few expressed genes, or are related to tumors/diseases/immunity, or have very similar gene sets (>60%) to other terms, we focused on 73 GO terms (**Table S8**). For each cell, the activities of the specific cellular functions represented by these GO terms were estimated by the AddModuleScore function of Seurat, which basically calculated the average Z-score transformed expression levels of all genes annotated by the GO term. All cells within a cluster were then combined to determine the average activity of the GO term for the cluster (**Figure 2B**).

### Transcriptome divergence among sub-CLTs

As for the divergence among sub-CLTs, estimation of pseudotime was conducted via Monocle^34^ with all cells on differentiating CLTs pooled together. After Principal Component Analysis of all cells from all samples combined, the transcriptomic divergence (*D*_T_) between any two cells is quantified by one minus Pearson’s Correlation Coefficient of the top 100 principal components. The developmental potential of an ancestor cell (an internal node on the CLT) was then calculated by the summed squared *D*_T_ of all pairs of its descendant cells. The reduction of developmental potential (*Δ*_DP_) during the growth of an internal node to its daughter nodes was calculated by the focal internal node’s *Δ*_DP_ subtracted by the summed *Δ*_DP_ of all its daughter nodes (**Figure 2D**). The statistical significance of an observed *Δ*_DP_ was estimated by contrasting the observation with its null distribution generated by random assignment of single-cell transcriptomes from hESC samples to the focal CLT (**Figure 2D** bottom right corner). We emphasized here that the null distribution should be estimated by the single-cell transcriptomes from the non-differentiating hESC sample, since using those from the differentiating CBRAD5 samples would introduce actual divergence into the null and thus lead to an underestimated statistical significance. It is also worth noting that this method is very similar to the commonly used nonparametric method of permutational multivariate analysis of variance (PERMANOVA^57^), except that Pearson’s correlation-based divergence replaces the distance-based divergence used in canonical PERMANOVA, as the correlation-based metric has consistently been shown to result in superior performance for single-cell transcriptomes^58,59^. We have also applied this PERMANOVA-based method to subsets of genes within the transcriptome. For example, only genes annotated with a specific GO term (**Table S8**) were used. A significant divergence for a specific GO term does not necessarily indicate a significant divergence in the whole transcriptome, since genes annotated with the GO term may have a small effect on the transcriptome as a whole. As a result, internal nodes with transcriptomic divergence do not necessarily represent a larger fraction than nodes with divergence on a specific GO term.

In order to perform a retrospective analysis of divergence progression, we need a normalized temporal scale that is comparable across samples. In theory, this scale could be derived from the mutation rate of the lineage barcode and/or the topological depth of a node (i.e., the number of nodes between the root and the focal node). Considering the variability in Cas9 editing efficiency over barcodes, as well as long inter-site deletions, we discarded the mutation rate-based scale. For the topological depth scale, due to both biological and experimental stochasticity, the reconstructed CLTs and their nodes have very different depths, despite the fact that they are supposed to correspond to the ten-day directed differentiation. Assuming that the internal nodes were evenly sampled on all root-to-tip paths throughout the CLT, the actual depth of a node should be reflected equally by its depth from the root and (indirectly) by the depth from the focal node to its descendent tips. Based on this logic, we defined the normalized depth of a node as *d* = (*d_r_*/*d_t_* + (1 − *d_s_*/*d_t_*))/2, where *d*_r_ is the focal node’s depth from root, *d*_t_ is the max depth found in the CLT, and the *d*_s_ is the max depth from the focal node to its descendent tips. Here, via division by *d*_t_, all depths were scaled from 0 to 1, with 0 being the root and 1 being the tips with maximal raw depth within the CLT.

### Transcriptional memory index

We followed previously proposed methods^29,39^ to calculate transcriptional memory index. In each cell type and for each gene expressed in >10% of cells of this type, the CV of the expression levels was calculated among all terminal cells of this type within a sub-CLT (containing at least two cells of this type). The minimal CV among all sub-CLTs, i.e. min(CV), was then used to represent the expression variability of the focal gene in this cell type. It was also calculated for each of 1,000 randomized CLTs created by reassigning all cells of the same type to a new lineage position that was originally occupied by the same cell type. These 1,000 min(*CV_Random_*) from randomized CLTs were averaged, i.e. mean (min (*CV_Random_*)), to yield a null expectation for the observed min(*CV*). Finally, the memory index was defined as *M* = (min(*CV*) - mean(min(*CV_Random_*))) / min(*CV*). Note that the final division by min(*CV*) is different from the previously defined memory index^29,39^, but allows comparisons between genes with very different baseline CVs or expression levels.

To test the hypothesized role of transcription factors in mediating transcriptional memory, we obtained lists of gene sets responsive to perturbations of individual transcription factors (“TF_Perturbations_Followed_By_Expression” in Enrichr^40^). The genes with highest memory indices (top 10% across all cell types) were assessed for enrichment in each of these TF-responsive gene sets using Enrichr ^40^. We reported (**Figure 3E**) the ‘‘combined score’’ calculated by Enrichr, which takes into account both the statistical significance and the magnitude of enrichment (combined score of enrichment *c* = log(*p*) * *o*, where *p* is the *P* value from Fisher’s exact test and *o* is the odds ratio of the enrichment^40^).

### Composition of terminal cell types compared among sub-CLTs and the full CLTs

To compare the terminal cell type composition of one sub-CLT with its expectation, we constructed a 2-by-*n* contingency table for the *n* cell types appearing in the entire CLT. The first row of the contingency table lists the observed count of terminal cells for each cell type within the focal sub-CLT. The second row of the table lists the expected count of each cell type as determined by the fractional cell type composition of the entire CLT multiplied by the size of the focal sub-CLT. We then calculated 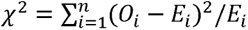 for the focal sub-CLT, where *O_i_* and *E_i_* are the observed and expected count for cell type *i*. Then *X*^2^ values from all sub-CLTs with roots of normalized depth < 0.7 (because roots closer to terminal cells produce sub-CLTs that are too small for meaningful statistics) were summed up to represent the diversity of cell type compositions among sub-CLTs (*x* axis of Figure 4A/B/C). In other words, a small summed *X*^2^ indicates uniform/stereotyped composition of cell types among sub-CLTs.

### Robustness of random versus stereotyped development

Without loss of generality, we defined a functional unit as consisting of four cell types, namely α, β, γ, and δ, in a 1:1:2:4 ratio. We simulated 1000 binary CLTs, each consisting of 1024 terminal cells (128 α cells, 128 β cells, 256 γ cells, 512 δ cells) generated through ten cell cycles, under two developmental models. The first “random” model randomly assigns the four types of cells onto the tips of the tree. A second “stereotyped” model strictly assigns α, β, γ, and δ cells in a 1:1:2:4 ratio onto each sub-CLT consisting of eight tips (three cell cycles). A predefined fraction (0.001, 0.005, 0.01, 0.05 or 0.1, as on *x* axis of **Figure 4E**) of the 2047 (1024 terminal and 1023 internal) cells were chosen and removed along with all their descendent cells to mimic random necrosis. Assuming sufficient cell migration to allow formation of the functional unit as long as there are enough terminal cells of the proper type, the robustness is thus quantified by the number of functional units that can be formed by all terminal cells surviving necrosis. A simple example shown in **Figure 4D**.

### Comparison and alignment of sub-CLTs by DELTA

To score the similarity of sub-CLTs assigned with different cell types, we transferred the expression distance (x) of cell node (terminal node) pair to the similar score (-log(x+1)) of cell node pair and calculation the similar score distribution of all cell node pairs from three differentiation samples and hESC sample. The cell node pair aligned with similarity less than 15 percentile cutoff of distribution was pruned, the pair with similarity more than 85 percentile cutoff of distribution was awarded from 0 based on the degree above the cutoff, lastly the pair with similarity between 15 percentile and 85 percentile was deduced from 0 based on the degree below the cutoff. The similar score of cell type pair was average score of cell node pairs labeled with same cell type pair. We implemented the alignment of sub-CLTs from any two CLT were based on the similar score of cell type pair. P value of any two sub-CLTs alignment was calculated with the order of similarity on the permutation distribution of 1000 times simulation CLT.

